# PremPRI: Predicting the Effects of Single Mutations on Protein-RNA Interactions

**DOI:** 10.1101/2020.04.07.029520

**Authors:** Ning Zhang, Haoyu Lu, Yuting Chen, Zefeng Zhu, Qing Yang, Shuqin Wang, Minghui Li

**Affiliations:** Center for Systems Biology, Department of Bioinformatics, School of Biology and Basic Medical Sciences, Soochow University, Suzhou 215123, China

## Abstract

Protein-RNA interactions are crucial for many cellular processes, such as protein synthesis and regulation of gene expression. Missense mutations that alter protein-RNA interaction may contribute to the pathogenesis of many diseases. Here we introduce a new computational method PremPRI, which predicts the effects of single mutations occurring in RNA binding proteins on the protein-RNA interactions by calculating the binding affinity changes quantitatively. The multiple linear regression scoring function of PremPRI is composed of 11 sequence- and structure-based features, and is parameterized on 248 mutations from 50 protein-RNA complexes. Our model shows a good agreement between calculated and experimental values of binding affinity changes with Pearson correlation coefficient of 0.72 and the corresponding root-mean-square error of 0.76 kcal mol^−1^, outperforming three other available methods. PremPRI can be used for finding functionally important variants, understanding the molecular mechanisms, and designing new protein-RNA interaction inhibitors. PremPRI is freely available at http://lilab.jysw.suda.edu.cn/research/PremPRI/.

## Introduction

The interactions between protein and RNA are crucial for many cellular processes, such as protein synthesis and regulation of gene expression [1–6]. Missense mutations occurring in these RNA binding proteins that alter protein-RNA interactions may cause significant deviation from the normal function of these proteins, potentially leading to various disorders including cancer [7, 8]. Indeed, several comprehensive studies of the structural nature of mutations in cancer and rare Mendelian diseases have shown that mutations located on binding interfaces may induce macromolecular interaction perturbations and play “driver” or “damaging” roles in many cancers and Mendelian diseases [9–16]. Quantifying the effects of missense mutations on specific protein-RNA interactions requires assessing the binding affinity changes upon introducing mutations, which can be accurately measured by traditional mutagenesis technologies. However, experimental methods used to measure binding affinity, such as surface plasmon resonance [17] and isothermal titration calorimetry [18], are costly and time-consuming. Therefore, developing reliable computational approaches provides an alternative way to investigate the effects of mutations on proteins and their interactions with other molecules on a large scale, and it will facilitate the identification of functionally important missense mutations and the discovery of the molecular mechanisms that cause diseases.

Many efforts have been made to computationally predict and model the effects of missense mutations on protein stability and protein-protein interactions [19–28]. However, predicting the impacts of mutations on protein–nucleic acid interactions has been more intractable and very few computational methods have been proposed [29–33]. One of the reasons that has hindered the development of methods is due to the complexity of nucleic acid chemistry and binding, which limited the availability of high-quality experimental data. Moreover, the interactions between protein-DNA and -RNA are different, which was clarified by a detailed comparison at the atomic contact level[34]. Recently, we developed a computational approach to estimate the impacts of missense mutations on protein-DNA interactions using molecular mechanics force fields and statistical potentials [30]. Peng et al. combined modified MM/PBSA based energy terms with additional knowledge-based terms to predict the protein-DNA binding affinity changes upon single mutations [31]. Pires et al. used graph-based signatures to model the effects of single mutations both on protein-DNA and -RNA binding [29]. The three methods mentioned above estimate the effects of mutations on the interactions by quantitatively calculating the changes in binding affinity. In addition, several classification computational approaches have been proposed for predicting the binding hot spots at protein-RNA binding interfaces [32, 33, 35]. Therefore, although some advancements have been made, the issue of predicting the effects of mutations on protein-RNA interactions is still at the initial stage.

To address this need, we introduced a new computational method, PremPRI, for characterizing the effects of single mutations on protein-RNA interactions by calculating the binding affinity changes quantitatively. PremPRI uses a novel multiple linear regression scoring function composed of 11 sequence- and structure-based features, and achieves significantly better performance than other predictors. PremPRI can be applied to many tasks, such as finding potential disease-causing and cancer driver missense mutations and understanding their molecular mechanisms, and designing inhibitors of protein-RNA interactions.

## Methods

### Experimental datasets used for training

First, we used ProNIT and dbAMEPNI databases to compile our training dataset. The ProNIT [36] includes experimentally measured values of changes in binding affinity (ΔΔ*G*) between proteins and nucleic acids upon single amino acid substitutions along with the experimentally determined 3D structures of protein-nucleic acid complexes. The dbAMEPNI [37], a recently proposed database, collected more experimentally measured binding affinity changes between protein and nucleic acid induced by alanine-scanning mutations compared to ProNIT. In order to construct an accurate and cleaned training dataset, we first removed the following complexes and mutations from the above databases, including the complexes with modified residues/nucleotides located at the protein-RNA binding interface, the complexes in which the protein length is less than 20 amino acids or RNA length is less than five nucleotides, and the mutations occurred at metal coordination sites. Then, to avoid inconsistencies between the RNA sequences used to measure binding affinity and the sequences presented in the 3D complex structures used to develop prediction model, we compared the sequence similarity between the sequences at the protein-RNA binding sites and the corresponding ones used to measure binding affinity. The RNA sequences in binding affinity measurements were either obtained from the ProNIT database or manually collected from the corresponding references. The entries with high sequence similarity (80%) were retained. In addition, there are three mutations with multiple experimental measurements of ΔΔ*G*. Since the differences between maximal and minimal ΔΔ*G* values for these cases are less than 1.0 kcal mol^−1^, the average value was used. As a result, 108 single mutations in 30 protein-RNA complexes from the ProNIT and dbAMEPNI databases were retained in our training dataset.

Secondly, 140 additional mutations obtained from the PrabHot benchmark and independent test sets were added into our training dataset [32], and they satisfied all above criteria. PrabHot is a classification computational method for identifying hot spots at the protein-RNA binding interfaces. Therefore, the final experimental dataset used to parameterize our PremPRI model includes 248 single mutations from 50 protein-RNA complexes (named as S248) (Table S1). The number of mutations for each protein-RNA complex is shown in Figure S1. In the S248 dataset, only 16 mutations have experimental pH values. Thus, the neutral pH was chosen at which the default charged states were assigned to the ionizable residues. We also compared our training dataset of S248 with them used for developing mCSM-NA [29] and PrabHot methods [32], and the details are shown in the Table S1.

### Structural optimization protocol

Three dimensional structures of protein-RNA complexes were obtained from the Protein Data Bank (PDB) [38]. The biological assembly 1 of crystal structure or the first model of NMR structure was used as the initial wild-type structure. Next we used BuildModel module of FoldX software package [20, 39] to produce mutant structures. Then VMD program [40] was applied to add missing heavy side-chain and hydrogen atoms to both wild-type and mutant structures using the topology file of CHARMM36 force field [41]. After that we carried out a 1000-step energy minimization for each complex in the gas phase during which the harmonic restraints with a force constant of 5 kcal mol^−1^ Å^−2^ were applied on the backbone atoms of all residues. The NAMD program v2.12 [42] and the CHARMM36 force field [41] were used to perform the energy minimization. A 12 Å cutoff distance for non-bonded interactions was applied to the molecular systems and lengths of hydrogen-containing bonds were constrained by the SHAKE algorithm [43]. The flowchart of structural optimization protocol is shown in Figure S2. The minimized wild-type and mutant protein-RNA complexes were used for the following calculations of energy features.

### The PremPRI model

The multiple linear regression scoring function of PremPRI is composed of 11 features and all of them have significant contribution to the quality of the model (p-value < 0.01, t-test). The p-value and the importance of each feature are presented in Table S2 and the description is illustrated below:

- ΔΔ*E*_*vdw*_ is the difference of Van der Waals interaction energies between mutant and wild type 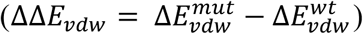. (Δ*E*_*vdw*_ is the difference of Van der Waals energies between a protein-RNA complex and each binding partner (partner1: protein; partner2: RNA), which is calculated using the ENERGY module of CHARMM program [44].
- ΔΔ*E*_*vdw.re*_ is the difference of Van der Waals repulsive energies between mutant and wild type. Here, the Van der Waals repulsive energy only counts the repulsion between the residue at the mutated site and the nucleotides.
- ΔΔ*E*_*elec*_ is the difference of electrostatic interaction energies between mutant and wild type 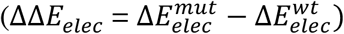. (ΔΔ*E*_*elec*_ is the electrostatic interaction energies between the residue at the mutated site and its contact residues/nucleotides. If any side-chain atom/base of a residue/nucleotide is located within 10 Å from any side-chain atom of the mutated site, we defined it as a contact residue/nucleotide. The calculation is carried out using the ENERGY module of CHARMM program.
- *N*_*inter*_ is the number of amino acids at the protein-RNA binding interface. If the solvent accessible surface area of a residue in the protein is more than that in the complex, we define it as the interface residue. The SASA module of CHARMM is used to calculate the solvent accessible surface area.
- *R*_*L/SA*_ is the ratio of protein length and its surface area. 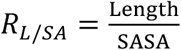, Length and SASA is the total number of residues and the solvent accessible surface area of unbound protein, respectively. The structure of unbound protein is extracted from the minimized wild-type complex structure.
- Closeness of the node of mutated site in the residue interaction network. It is defined as:

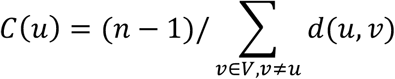

where *d*(*u*, *v*) is the shortest-path distance between the node u of mutated site and any node *v*. V is the set of all nodes and n is the number of nodes in the residue interaction network. The shortest-path distance between two nodes refers to the minimum number of nodes that reach from one node to the other [45]. Cα atom of a residue is considered as a node. If the distance between two Cα atoms is less than 6 Å, we define them as having a direct interaction. Closeness is calculated using the Python package NetworkX [46].
- Δ*SA* is the difference of solvent accessible surface areas between mutant and wild type 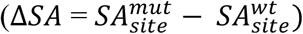. *SA*_*site*_ is the solvent accessible surface area of the residue at the mutated site in the unbound protein that is extracted from the minimized complex structure.
- 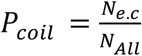, *N*_*e.c*_ and *N*_*All*_ are the number of exposed residues in the coil conformation and all residues in the mutated protein chain, respectively. Secondary structure elements other than α-helices and β-strands are defined as coil, which are assigned by DSSP program [47]. If the ratio of the solvent accessible surface area of a residue in the complex and in solvent is more than 0.25 [48], we defined it as the exposed residue.
- *ΔOMH* is the difference of hydrophobicity scale between mutant and wild-type residue type. The hydrophobicity scale (OMH) for each type of amino acid was derived by considering the observed frequency of amino acid replacements among thousands of related structures, which was taken from the study of [49] directly.
- Δ*P*_FWY_ and 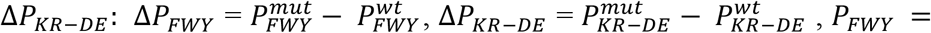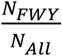 and 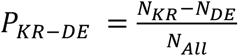 and *N*_*All*_ are the number of aromatic (F, W and Y), positively charged (K and R), negatively charged (D and E) and all amino acids in the mutated protein chain, respectively.

In addition, we also tested the performance of several other popular machine learning algorithms, including Random Forest (RF), Back Propagation Neural Network (BPNN), Support Vector Machine (SVM) and eXtreme Gradient Boosting (XGBoost), and the results shown in Table S3 indicate that the multiple linear regression algorithm presents the best performance.

### Statistical analysis and evaluation of performance

Pearson correlation coefficient (R) and Root Mean Square Error (RMSE) are used to measure the agreement between experimentally determined and predicted values of binding affinity change. All correlation coefficients presented in the paper are significantly different from zero (p-value < 0.01, t-test). RMSE (kcal mol^−1^) is the standard deviation of the residuals (prediction errors). Hittner2003 test [50, 51] is used for comparing whether the difference in correlation coefficients between PremPRI and other methods is significant. The changes in the area under the receiver operating characteristics curve (AUC of ROC) are tested by Delong test [52]. All tests are implemented in R.

Receiver Operating Characteristics and precision-recall analyses were performed to quantify the performance of different methods in distinguishing mutations highly decreasing binding affinity (ΔΔ*G*_*exp*_ ≥ 1 kcal mol^−1^) from others. True positive rate (TPR) and false positive rate (FPR) is defined as TPR=TP/(TP + FN) and FPR=FP/(FP+TN) (TP: true positive; TN: true negative; FP: false positive; FN: false negative) respectively. In addition, the maximal Matthews correlation coefficient (MCC) value is reported for each method by calculating the MCC across a range of thresholds:

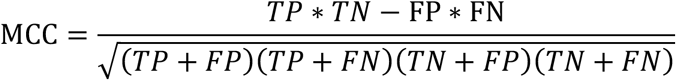

## RESULTS and DISCUSSION

### Multiple linear regression model of PremPRI

The PremPRI model is built using multiple linear regression algorithm and composed of ten structure- and one sequence-based features. The p-value and importance for each feature are shown in Table S2 that indicate all of the features contribute significantly to the model. To investigate the linear associations between different features, we performed multicollinearity analysis. Table S4 presents that the variance inflation factor (VIF) of each feature is less than three, indicating low collinear relationships among 11 independent variables in PremPRI model.

The performance of PremPRI trained and tested on the S248 set is shown in Figure 1a and Table 1. The Pearson correlation coefficient between experimental and calculated binding affinity change is 0.72 and the corresponding RMSE and slope is 0.76 kcal mol^−1^ and 1.00 respectively.

**Table 1.**
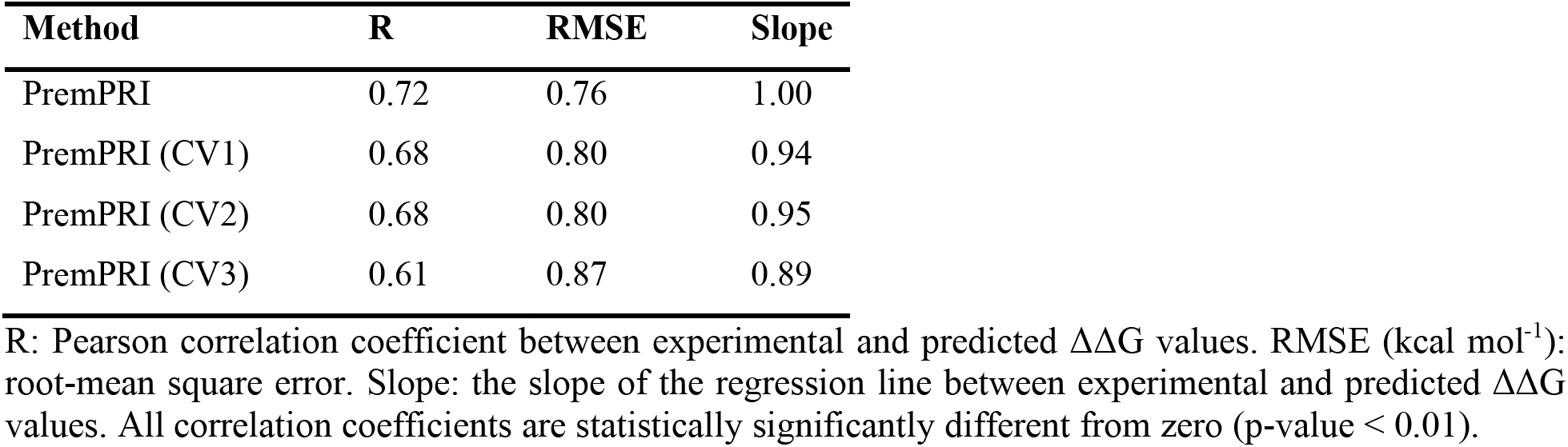
The performance for PremPRI trained and tested on S248 dataset and performing three types of cross-validation.

| Method | R | RMSE | Slope |
| --- | --- | --- | --- |
| PremPRI | 0.72 | 0.76 | 1.00 |
| PremPRI (CV1) | 0.68 | 0.80 | 0.94 |
| PremPRI (CV2) | 0.68 | 0.80 | 0.95 |
| PremPRI (CV3) | 0.61 | 0.87 | 0.89 |

R: Pearson correlation coefficient between experimental and predicted $\Delta\Delta G$ values. RMSE (kcal mol<sup>-1</sup>): root-mean square error. Slope: the slope of the regression line between experimental and predicted $\Delta\Delta G$ values. All correlation coefficients are statistically significantly different from zero (p-value < 0.01).

**Figure 1.**
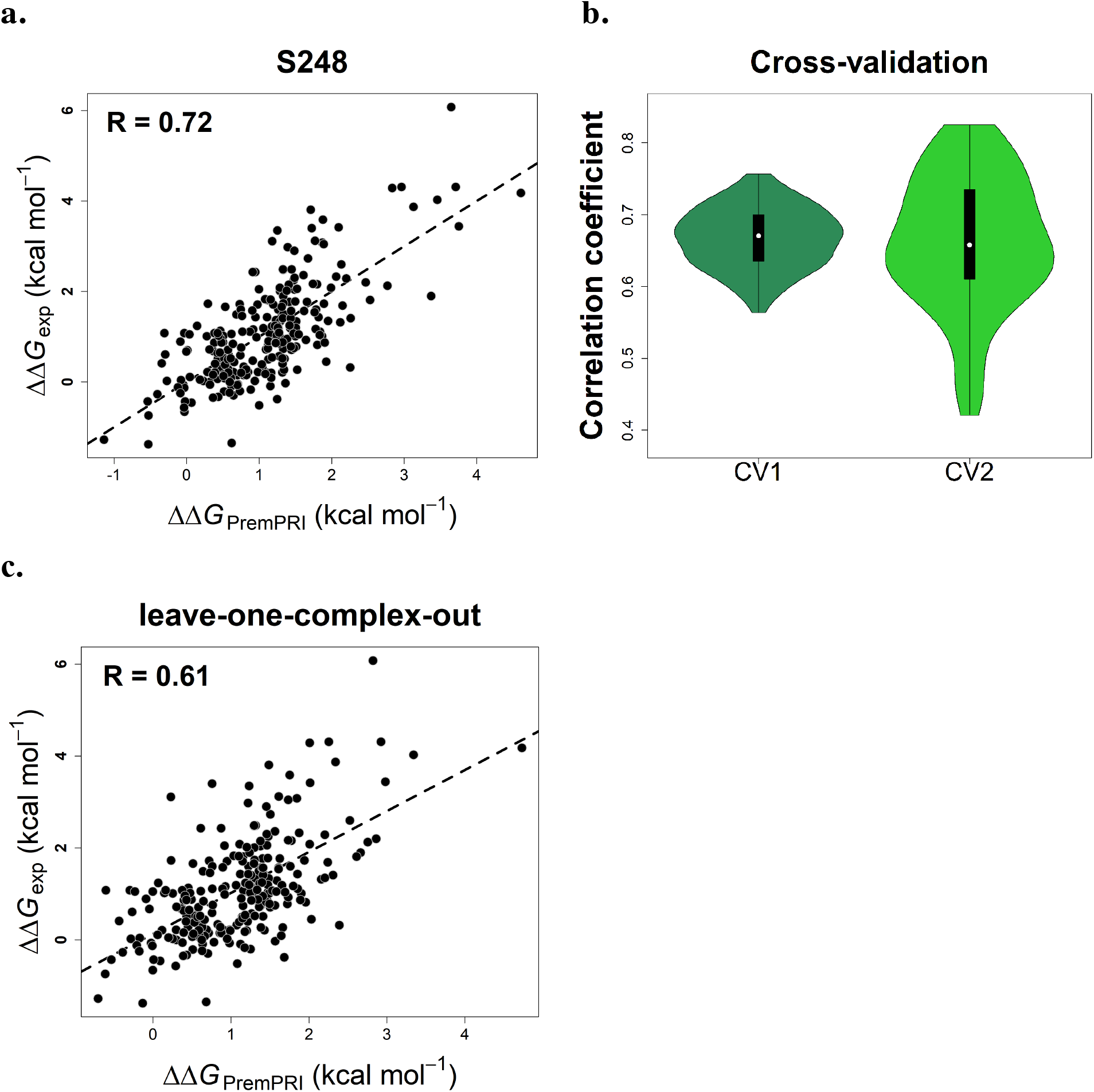
Pearson correlation coefficients between experimental and calculated changes in binding affinity for PremPRI trained and tested on S248 dataset (a), using two types of cross-validation (CV1 and CV2) (b) and performing leave-one-complex-out validation (CV3) (c), respectively.

PremPRI takes about four minutes to perform the calculation for a single mutation in a protein-RNA complex with ~ 330 residues and ~ 70 nucleotides, and requires 30 seconds for each additional mutation introduced in the same complex.

### Performance on three types of cross-validation

Overfitting is one of the major problems in machine learning. In order to check whether our method has this issue, we performed three types of cross-validation (CV1, CV2 and CV3). In the CV1 and CV2, we randomly chose 50% and 80% of all mutations from the S248 dataset to train the model, respectively, and used the remaining mutations for blind testing; both procedures were repeated 100 times. In the CV3, the model was trained and tested using non-overlapping sets of protein-RNA complexes. Namely, we left one complex and its mutations out as the test set and trained the model using the rest of the complexes/mutations (leave-one-complex-out validation); this process was repeated for each complex.

The correlation coefficients for 100 times cross-validation of CV1 and CV2 are shown in Figure 1b, and the R of each round is greater than 0.42. The average values of R and RMSE for 100 times CV1 and CV2 are 0.68 and 0.80 kcal mol^−1^ respectively (Table 1). In the leave-one-complex-out validation (CV3), the correlation coefficient reaches to 0.61 (Fig. 1c) and RMSE = 0.87 kcal mol^−1^ (Table 1). Moreover, PremPRI does not present bias to the different categories of mutations, including alanine-scanning and non-alanine-scanning mutations, interfacial and non-interfacial mutations, and the mutations from protein-single stranded RNA complexes (Protein-ssRNA) and protein-double stranded RNA complexes (Protein-dsRNA) (Table S1 and Table S5). The performance for each category remains relatively high in CV3 with R of ~ 0.60 (Table S5). In addition, analysis of the variation of weighting coefficient for each feature in the three types of cross-validation further illustrates that the PremPRI model does not overfit on its training set and all features contribute significantly to the energy function (Table S6).

### Comparison with other methods

We compared our method with three other available computational methods, mCSM-NA [29], PrabHot [32] and FoldX5.0 [39], developed for predicting the effects of mutations on protein-RNA interactions. mCSM-NA calculates the changes in protein-DNA/RNA binding affinity using graph-based signatures. PrabHot is a classification method and uses a combination of sequence-, structure- and residue-interaction-network-based features to identify hot spots at protein-RNA binding interfaces. FoldX5.0 calculates the binding affinity changes using an empirical energy function and is parametrized on experimentally determined unfolding free energy changes. All of them are machine learning approaches and the training datasets for parameterizing mCSM-NA and PrabHot are shown in the Table S1. The number of common mutations between S248 and the training datasets of mCSM-NA and PrabHot is 16 and 92 respectively. We did not compare S248 with the training dataset of FoldX5.0 since it is composed of unfolding free energy changes.

We applied all three methods on the S248 dataset, and the correlation coefficients are 0.20 (RMSE = 1.57 kcal mol^−1^) and 0.24 (RMSE = 5.41 kcal mol^−1^) for FoldX and mCSM-NA respectively (Table 2). Moreover, we performed ROC and precision-recall analyses in order to estimate the performance of different methods to predict the highly decreasing mutations (ΔΔ*G*_*exp*_ ≥ 1 kcal mol^−1^, the same cutoff was used in PrabHot to define the hot spots), and the results are also shown in the Table 2. For PremPRI, the leave-one-complex-out validation results are used. Although it is not the completely identical comparison, the significantly large differences of the values of R, RMSE, AUC-ROC, AUC-PR and MCC between PremPRI (CV3) and the other methods can prove the better performance for our method. In addition, our method performs well for both interfacial and non-interfacial mutations (Fig. 2b), and only statistically significant correlation coefficient is observed for mCSM-NA to predict interfacial mutations.

**Table 2.**
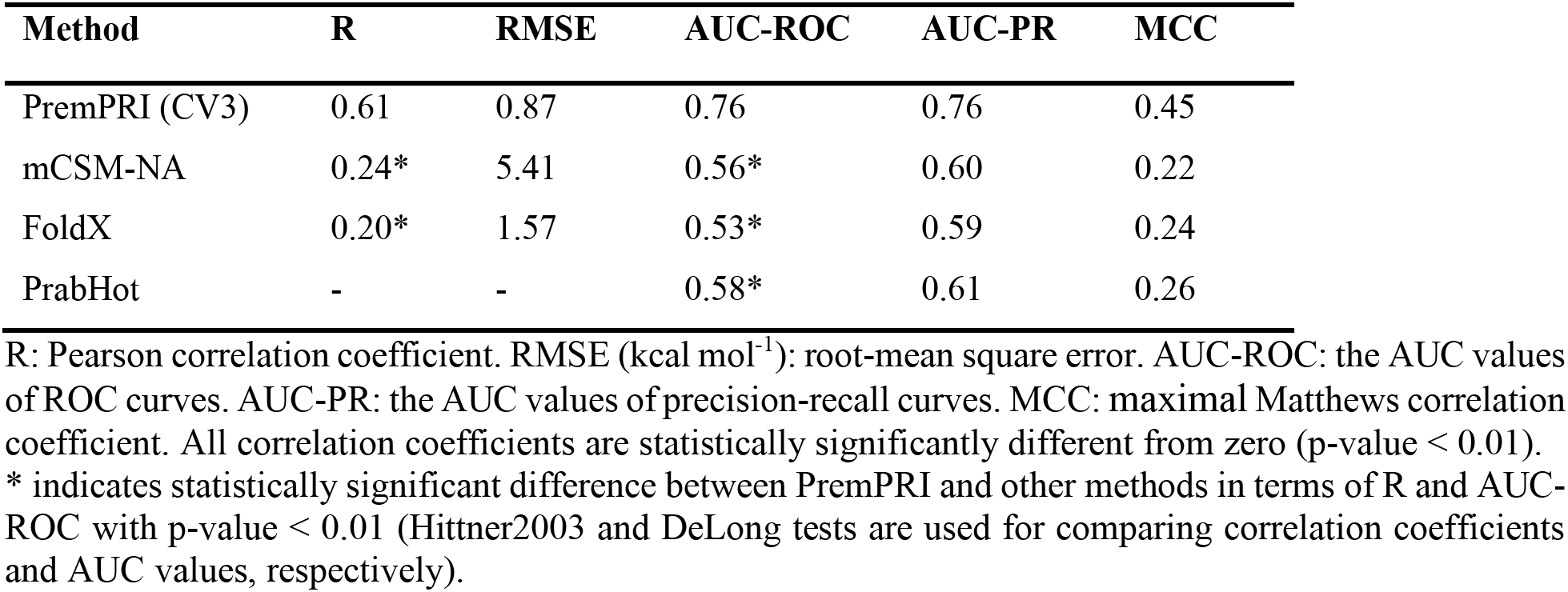
Comparison of methods’ performances on S248 dataset.

**Figure 2.**
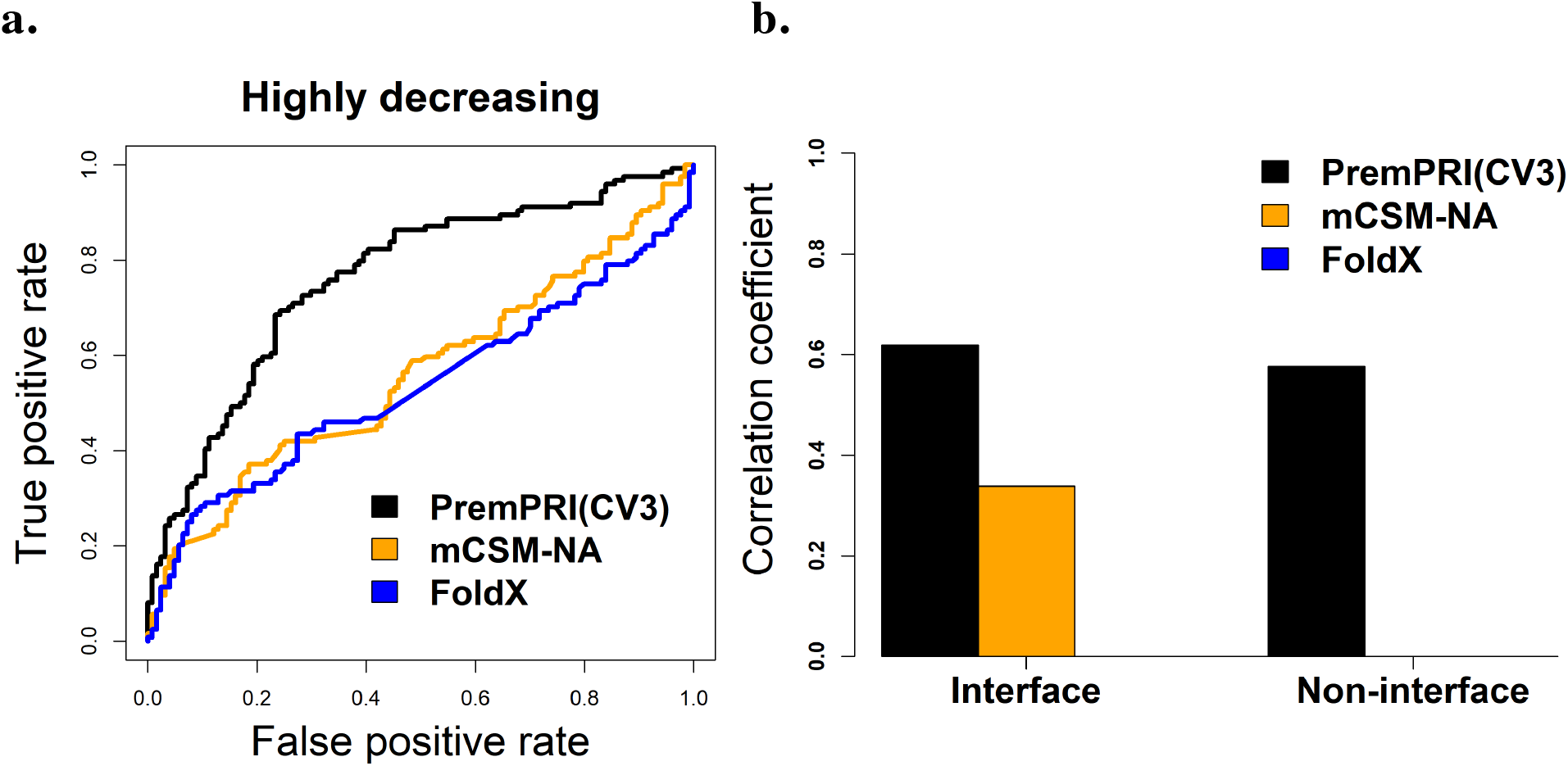
Performance of three methods of PremPRI, mCSM-NA and FoldX applied on the S248 dataset. The leave-one-complex-out validation (CV3) results of PremPRI are used. (a) ROC curves for predicting highly decreasing mutations. The number of highly decreasing mutations (ΔΔ*G*_*exp*_ ≥ 1 kcal mol^−1^) in S248 is 124. (b) Pearson correlation coefficients between predicted and experimental ΔΔ*G* for mutations located at protein-RNA binding interface and non-interface. Only correlation coefficients that are significantly different from zero are shown in the figure (p-value < 0.01, t-test).

Furthermore, we applied all methods on a test case that includes three single mutations occurring in the highly conserved triad Thr-Met-Gly of Ribosomal protein L1 [53]. All three mutations lower the protein-RNA binding largely (The experimental values of ΔΔ*G*_*exp*_ are presented in Table S7). Crystal structure of Ribosomal protein L1 from Thermus thermophilus (TthL1) in complex with a specific 80 nt fragment of 23S rRNA (PDB ID: 3U4M) is used to perform the calculations. Our training dataset of S248 includes one mutation of T217A from this complex of 3U4M, which was excluded from the training dataset when testing on this case. The predictions shown in the Table S7 indicate that the PremPRI has the best performance and predicts all three mutations as highly decreasing mutations.

## ONLINE WEBSERVER

### Input

The PremPRI webserver requires the 3D structure of a protein-RNA complex that can be retrieved from the Protein Data Bank with the PDB code input or provided by a file with the atomic coordinates uploaded by the user (Fig. S3a). In either case, the structure file must include at least two chains, one is protein and the other is RNA. After the structure is retrieved correctly, the server will display a 3D view of the complex structure and list the corresponding protein or RNA name for each chain (Fig. S3b). At the second step, two interaction partners should be defined, and one or multiple chains can be assigned to each partner. Only assigned chains will be considered during the prediction. At the third step of selecting mutations, three options are provided allowing users to do large-scale mutational scanning (see Fig. S3c). In the option of “Specify One or More Mutations Manually”, the user can submit the specified mutations and visualize each mutated site in the protein-RNA complex structure. “Alanine Scanning for Each Chain” option is used to perform alanine scanning for each protein chain. The “Upload Mutation List” option allows users to submit a list of mutations specified in the uploaded file.

### Output

For each single mutation in a protein-RNA complex, the PremPRI server provides: ΔΔG (kcal mol^−1^), predicted binding affinity change, and positive and negative signs correspond to the mutations decreasing and increasing binding affinity respectively; Interface (yes/no), shows whether the mutation occurs at the protein-RNA binding interface. When a residue’s solvent accessible surface area in the complex is lower than in the unbound partner, it is defined as located at the interface. Furthermore, for each mutation, PremPRI provides an interactive 3D viewer which shows the non-covalent interactions between the residue in the mutated site and its adjacent residues/nucleotides, generated by Arpeggio [54]. The minimized wild-type and mutant complex structures are used to show the interactions. An example is provided in Figure S4.

## ACKNOWLEDGEMENT

This research was supported by the National Natural Science Foundation of China (Grant No. 31701136), Natural Science Foundation of Jiangsu Province, China (Grant No. BK20170335), and the Priority Academic Program Development of Jiangsu Higher Education Institutions.

