## Supplementary Materials for "PremPRI: Predicting the Effects of Single Mutations on Protein-RNA Interactions"

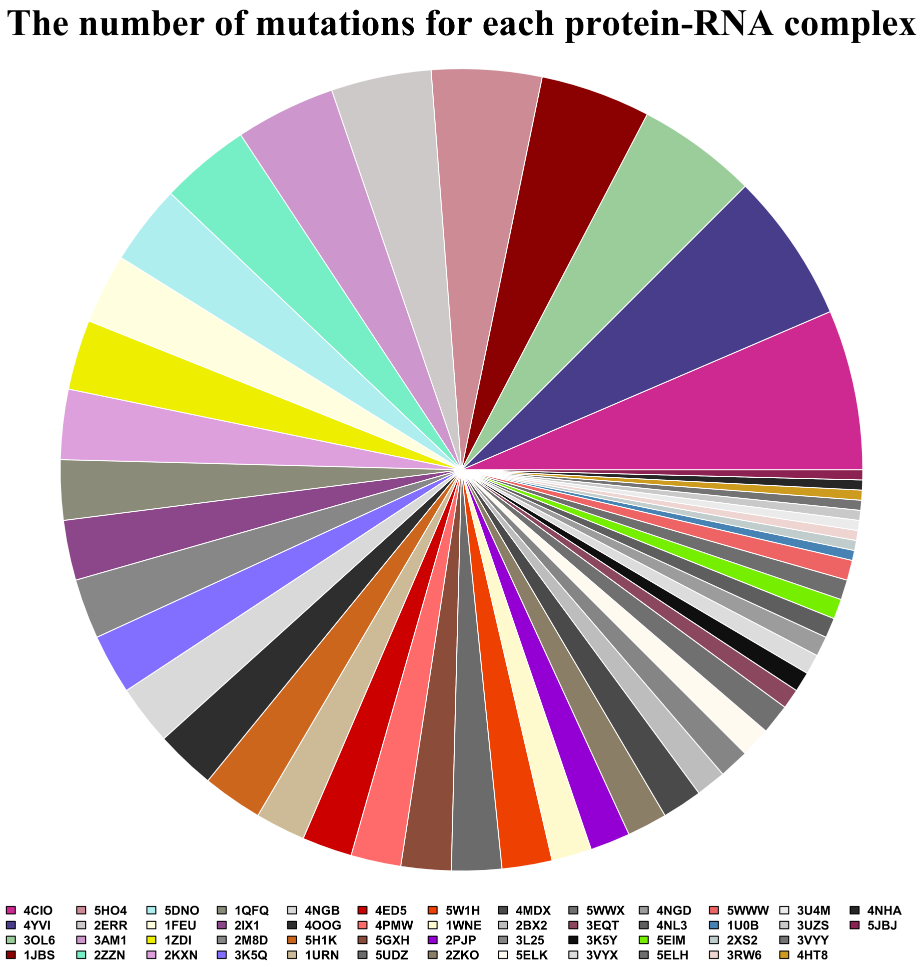


**Figure S1.** The number of mutations for each protein-RNA complex in S248 dataset.

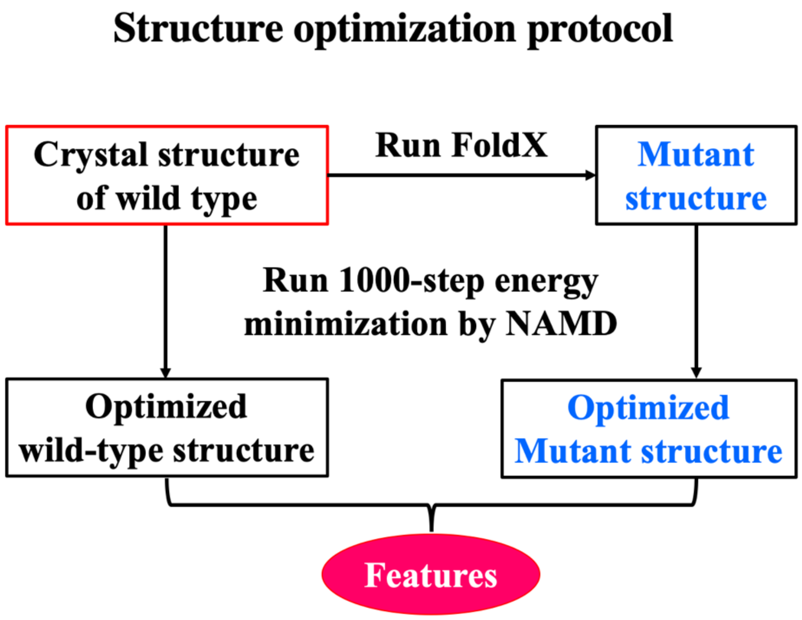


**Figure S2.** The flowchart of structure optimization protocol.

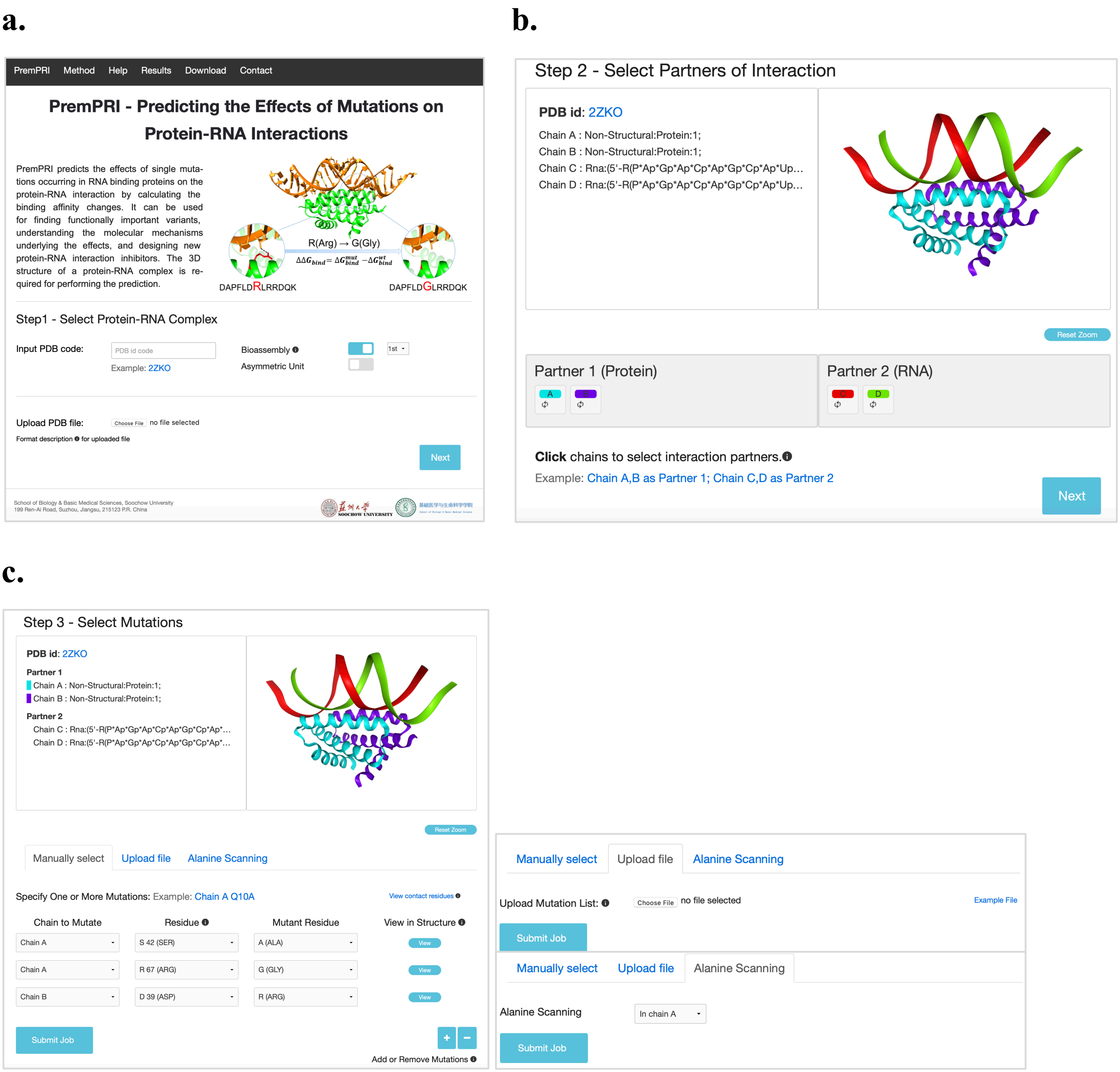


**Figure S3.** (a) The entry page of PremPRI server. (b) The second step for selecting interaction partners. (c) The third step for selecting mutations and three options are provided: "Specify One or More Mutations Manually", "Upload Mutation List" and "Alanine Scanning for Each Chain".

**a.**

**Job id:** 2020050504060173705807592

- Summary

| PDB ID | Protein | RNA | Number of mutations | Start time (EST) | Processing time | Results |
|---|---|---|---|---|---|---|
| 2ZKO | A, B | C, D | 3 | 2020-05-04 23:09 | 5 min | Download |

- Results

| # | Mutated Chain | Mutation | ΔΔG | Interface? | Structure |
|---|---|---|---|---|---|
| 1 | A | S42A | **1.5** | Yes | Explore |
| 2 | A | R67G | **1.03** | No | Explore |
| 3 | B | D39R | **2.91** | Yes | Explore |

Click

**b.**

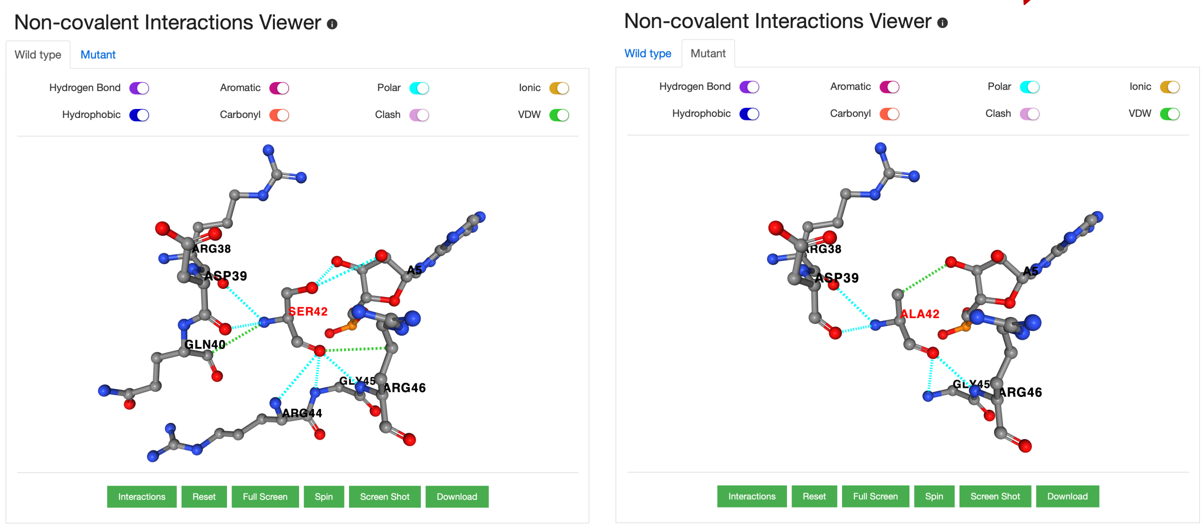


**Figure S4.** (a) The final results. "Process time" refers to the running time of a job without counting the waiting time in the queue. (b) Interactive 3D viewer showing the non-covalent interactions between the mutated site in the NS1 protein of human influenza virus A (PDB ID: 2ZKO, mutation: S42A) and its adjacent residues/nucleotides in the wild-type (left) and mutant (right) complex respectively, generated by Arpeggio.

**Table S1. Experimental datasets used for training methods of PremPRI, mCSM-NA and PrabHot.**

| Dataset | # of mutations | # of complexes | Description |
|---|---|---|---|
| S248 | 248 | 50 | training set of PremPRI |
| S264 | 264(67) | 33(5) | training set of mCSM-NA including mutations from both protein-DNA and -RNA complexes; bracket: the number of mutations from protein-RNA complexes |
| S151 | 151 | 32 | training set of PrabHot, classification method |
| S16 | 16 | 2 | overlap mutations between S248 and S264 |
| S92 | 92 | 21 | overlap mutations between S248 and S151 |

**Different categories for mutations in S248**

| Category | # of mutations | # of complexes | Description |
|---|---|---|---|
| Alanine-scanning | 213 | 50 | substitutions of residues into alanine |
| Non-alanine-scanning | 35 | 13 | substitutions of residues into non-alanine |
| Interface | 154 | 45 | mutations occur at protein-RNA binding interface |
| Non-interface | 94 | 31 | mutations do not occur at binding interface |
| Protein-ssRNA | 122 | 24 | mutations occur in protein-single stranded RNA complexes |
| Protein-dsRNA | 126 | 26 | mutations occur in protein-double stranded RNA complexes |

**Table S2. The p-value and importance of each feature in multiple linear regression scoring function of PremPRI.**

| Feature | P-value | Importance |
|---|---|---|
| $\Delta \boldsymbol{P}_{\boldsymbol{FWY}}$ | 1.40e-13 | 0.52 |
| $\boldsymbol{N}_{\boldsymbol{inter}}$ | 7.24e-07 | 0.30 |
| $\boldsymbol{Closeness}$ | 6.90e-06 | 0.30 |
| $\boldsymbol{R}_{\boldsymbol{L/SA}}$ | 1.23e-05 | 0.29 |
| $\Delta\Delta \boldsymbol{E}_{\boldsymbol{vdw.re}}$ | 1.18e-05 | 0.28 |
| $\boldsymbol{P}_{\boldsymbol{coil}}$ | 2.07e-05 | 0.26 |
| $\Delta\Delta \boldsymbol{E}_{\boldsymbol{elec}}$ | 1.49e-03 | 0.20 |
| $\Delta \boldsymbol{SA}$ | 7.78e-04 | 0.19 |
| $\Delta \boldsymbol{P}_{\boldsymbol{KR-DE}}$ | 5.13e-04 | 0.19 |
| $\Delta \boldsymbol{OMH}$ | 3.27e-03 | 0.19 |
| $\Delta\Delta \boldsymbol{E}_{\boldsymbol{vdw}}$ | 3.59e-03 | 0.15 |

Standardized coefficients are used for describing the importance.

**Table S3. The performance using multiple linear regression (MLR), Random Forest (RF), Back Propagation Neural Network (BPNN), Support Vector Machine (SVM) and eXtreme Gradient Boosting (XGBoost) algorithms to build PremPRI model, respectively.**

| Algorithm | Method | R | RMSE | Slope |
|---|---|---|---|---|
| **MLR** | PremPRI | 0.72 | 0.76 | 1.00 |
| | PremPRI (CV3) | 0.61 | 0.87 | 0.89 |
| **RF** | PremPRI | 0.70 | 0.79 | 1.21 |
| | PremPRI (CV3) | 0.46 | 0.98 | 1.15 |
| **BPNN** | PremPRI | 0.82 | 0.63 | 1.01 |
| | PremPRI (CV3) | 0.46 | 0.99 | 0.72 |
| **SVM** | PremPRI | 0.85 | 0.61 | 1.25 |
| | PremPRI (CV3) | 0.42 | 1.00 | 0.89 |
| **XGBoost** | PremPRI | 0.99 | 0.14 | 1.09 |
| | PremPRI (CV3) | 0.40 | 1.00 | 0.95 |

PremPRI: trained and tested on S248 dataset; PremPRI (CV3): leave-one-complex-out validation results.

R: Pearson correlation coefficient. RMSE (kcal $mol^{-1}$): root-mean-square error. Slope: the slope of the regression line between experimental and predicted $\Delta\Delta G$ values. All presented correlation coefficients are statistically significantly different from zero (p-value << 0.01, t-test).

**Table S4. Variance inflation factor (VIF) of each feature in PremPRI model.**

| $\Delta\Delta E_{vdw}$ | $\Delta P_{KR-DE}$ | $\Delta SA$ | $N_{inter}$ | $P_{coil}$ | $\Delta\Delta E_{elec}$ | $\Delta\Delta E_{vdw.re}$ | *Closeness* | $R_{L/SA}$ | $\Delta OMH$ | $\Delta P_{FWY}$ |
|---|---|---|---|---|---|---|---|---|---|---|
| 1.25 | 1.41 | 1.61 | 1.69 | 1.81 | 1.85 | 1.97 | 2.07 | 2.12 | 2.13 | 2.20 |

**Table S5. PremPRI performance for different categories of mutations.**

| Mutation category | Method | R | RMSE | Slope |
|---|---|---|---|---|
| **Alanine-scanning** | PremPRI | 0.71 | 0.74 | 1.01 |
| | PremPRI (CV3) | 0.61 | 0.83 | 0.89 |
| **Non-alanine-scanning** | PremPRI | 0.78 | 0.85 | 0.98 |
| | PremPRI (CV3) | 0.60 | 1.08 | 0.90 |
| **Interface** | PremPRI | 0.75 | 0.78 | 1.05 |
| | PremPRI (CV3) | 0.62 | 0.93 | 0.93 |
| **Non-interface** | PremPRI | 0.65 | 0.71 | 0.86 |
| | PremPRI (CV3) | 0.58 | 0.77 | 0.76 |
| **Protein-ssRNA** | PremPRI | 0.76 | 0.82 | 1.08 |
| | PremPRI (CV3) | 0.61 | 0.98 | 0.99 |
| **Protein-dsRNA** | PremPRI | 0.64 | 0.69 | 0.84 |
| | PremPRI (CV3) | 0.59 | 0.75 | 0.74 |

PremPRI: trained and tested on S248 dataset; PremPRI (CV3): leave-one-complex-out validation results.

R: Pearson correlation coefficient. RMSE (kcal $mol^{-1}$): root-mean-square error. Slope: the slope of the regression line between experimental and predicted $\Delta\Delta G$ values. All presented correlation coefficients are statistically significantly different from zero (p-value << 0.01, t-test).

**Table S6. Average weighting coefficient and the corresponding standard deviation (in bracket) for each feature in three types of cross-validation. The weighting coefficient in the PremPRI model is presented for the comparison.**

| | CV1 | CV2 | CV3 | PremPRI |
|---|---|---|---|---|
| **Intercept** | -0.33(0.54) | -0.39(0.30) | -0.41(0.14) | -0.41 |
| $\Delta\Delta\boldsymbol{E}_{\boldsymbol{elec}}$ | 1.07e-03(3.66e-04) | 1.16e-03(1.72e-04) | 1.14e-03(3.90e-05) | 1.14e-03 |
| $\Delta\boldsymbol{SA}$ | 5.47e-03(1.70e-03) | 5.56e-03(9.66e-04) | 5.52e-03(3.34e-04) | 5.54e-03 |
| $\boldsymbol{N}_{\boldsymbol{inter}}$ | -9.55e-03(2.01e-03) | -9.27e-03(7.24e-04) | -9.21e-03(4.41e-04) | -9.20e-03 |
| $\Delta\Delta\boldsymbol{E}_{\boldsymbol{vdw}}$ | 0.02(6.90e-03) | 0.02(3.81e-03) | 0.02(1.20e-03) | 0.02 |
| $\Delta\Delta\boldsymbol{E}_{\boldsymbol{vdw.re}}$ | -0.11(2.57e-02) | -0.11(1.19e-02) | -0.11(5.31e-03) | -0.11 |
| $\Delta\boldsymbol{OMH}$ | 0.21(0.06) | 0.21(0.03) | 0.21(1.42e-02) | 0.21 |
| $\boldsymbol{P}_{\boldsymbol{coil}}$ | -5.95(1.68) | -5.71(0.81) | -5.66(0.44) | -5.67 |
| $\boldsymbol{Closeness}$ | 5.59(1.33) | 5.66(0.77) | 5.77(0.31) | 5.77 |
| $\Delta\boldsymbol{P}_{\boldsymbol{KR-DE}}$ | -31.65(16.34) | -32.70(7.45) | -33.34(3.73) | -33.48 |
| $\boldsymbol{R}_{\boldsymbol{L/SA}}$ | 83.93(17.89) | 83.01(10.69) | 83.53(4.26) | 83.51 |
| $\Delta\boldsymbol{P}_{\boldsymbol{FWY}}$ | -218.13(31.43) | -216.40(15.44) | -215.51(9.57) | -216.23 |

**Table S7. Comparison of methods' performances on three mutations from TthL1–RNA complex.** $\Delta\Delta G_{exp}$ and $\Delta\Delta G_{pred}$ are experimentally determined and predicted binding affinity change (in kcal $mol^{-1}$), respectively.

| **Mutation** | **$\Delta\Delta G_{exp}$** | **PremPRI** | **mCSM-NA** | **FoldX** | **PrabHot** |
|---|---|---|---|---|---|
| T217A | 2.49 | 1.32 | -1.18 | -1.22 | hotspot |
| T217V | 3.61 | 1.87 | 1.20 | 0.12 | hotspot |
| M218L | 6.58 | 1.67 | 1.59 | 0.13 | hotspot |
| G219V | 5.35 | 1.94 | -1.53 | 0.24 | non-hotspot |

Our training dataset of S248 includes one mutation of T217A from this complex, which was excluded from the training dataset when testing on this case.
